# The genetic prehistory of the Andean highlands 7,000 Years BP though European contact

**DOI:** 10.1101/381905

**Authors:** John Lindo, Randall Haas, Courtney Hofman, Mario Apata, Mauricio Moraga, Ricardo Verdugo, James T. Watson, Carlos Viviano Llave, David Witonsky, Enrique Vargas Pacheco, Mercedes Villena, Rudy Soria, Cynthia Beall, Christina Warinner, John Novembre, Mark Aldenderfer, Anna Di Rienzo

## Abstract

The peopling of the Andean highlands above 2500m in elevation was a complex process that included cultural, biological and genetic adaptations. Here we present a time series of ancient whole genomes from the Andes of Peru, dating back to 7,000 calendar years before present (BP), and compare them to 64 new genome-wide genetic variation datasets from both high and lowland populations. We infer three significant features: a split between low and high elevation populations that occurred between 9200-8200 BP; a population collapse after European contact that is significantly more severe in South American lowlanders than in highland populations; and evidence for positive selection at genetic loci related to starch digestion and plausibly pathogen resistance after European contact. Importantly, we do not find selective sweep signals related to known components of the human hypoxia response, which may suggest more complex modes of genetic adaptation to high altitude.

**One Sentence Summary:** Ancient DNA from the Andes reveals a complex picture of human adaptation from early settlement to the colonial period.

---

The Andean highlands of South America have long been considered a natural laboratory for the study of genetic adaptation of humans (*1*), yet the genetics of Andean highland populations remain poorly understood. People likely entered the highlands shortly after their arrival on the continent (*2, 3*), and while some have argued that humans lived permanently in the Central Andean highlands by 12,000 BP (*4*), other research indicates that permanent occupation began between 9500-9000 BP (*5–7*). Regardless of when the peopling of high elevation environments began, selective pressure on the human genome was likely strong due to challenging environmental factors but also social processes such as the intensification of subsistence resources and residential sedentism (*8*), which in turn promoted the development of agricultural economies, social inequality, and relatively high population densities across much of the highlands. European contact initiated an array of economic, social, and pathogenic changes (*9,10*). Although it is known that the peoples of the Andean highlands experienced population contraction after contact (*11*), its extent is debated from both archaeological and ethnohistorical perspectives, especially concerning the size of the indigenous populations at initial contact (*12, 13*).

To address hypotheses regarding the population history of Andean highlanders and their genetic adaptations, we collected a time series of ancient whole genomes from individuals in the Lake Titicaca region of Peru (Fig. S1, Table 1). The series represents three different cultural periods, which include individuals from (A) Rio Uncallane, a series of cave crevice tombs dating to ~1,800 BP and used by fully sedentary agriculturists, (B) Kaillachuro, a ~3,800 year old site marked by the transition from mobile foraging to agropastoralism and residential sedentism, and (C) Soro Mik’aya Patjxa (SMP), an 8000-6500 year old site inhabited by residentially mobile hunter-gatherers (*14*). We then compare the genomes of these ancient individuals to 25 new genomes and 39 new genome-wide SNP datasets generated from two modern indigenous populations: the Aymara of highland Bolivia and the Huilliche-Pehuenche of coastal lowland Chile. The Aymara are an agropastoral people who have occupied the Titicaca basin for at least 2,000 years (*15*). The Huilliche-Pehuenche are traditionally hunter-gatherers from the southern coastal forest of Chile (*16*).

**Table 1.** Ancient Sample Sequencing results

| **Table 1. Ancient Sample Sequencing results** |  |  |  |  |  |  |  |  |  |  |
| --- | --- | --- | --- | --- | --- | --- | --- | --- | --- | --- |
| ID | <sup>14</sup>C BP | mode<br>cal. BP<sup>2</sup> | 95% cal.<br>BP<sup>4</sup> | mtDNA<br>Haplo-<br>group | Sex<sup>1</sup> | Site | Endogenous<br>DNA | Mapped<br>Reads | Average<br>Read<br>Depth | Whole<br>Genome<br>Coverage |
| IL2 | Not dated | NA | NA | D1 | XX | Rio<br>Uncallane | 38.6% | 193316050 | 6.51 | 0.99 |
| IL3 | 1830±20 | 1711 | 1783-1616 | B2 | XX | Rio<br>Uncallane | 29.9% | 175348009 | 5.89 | 0.92 |
| IL4 | 1630±20 | 1514 | 1529-1426 | C1b | XY | Rio<br>Uncallane | 17.1% | 67544346 | 2.31 | 0.75 |
| IL5 | 1805±20 | 1703 | 1722-1611 | C1c | XX | Rio<br>Uncallane | 25.2% | 88991036 | 3.34 | 0.94 |
| IL7 | 1885±20 | 1806 | 1825-1730 | C1b | XY | Rio<br>Uncallane | 44.9% | 184150720 | 5.62 | 0.98 |
| K1 | Insufficient<br>collagen | NA | NA | C1b | XX | Kaillachuro | 2.60% | 12815292 | 1.44 | 0.26 |
| SMP5 | 6030±20 | 6822 | 6884-6757 | C1c | XY | Soro<br>Mik'aya<br>Patjxa | 3.15% | 3655632 | 3.92 | 0.08 |
| <sup>1</sup>Sex was determined from sequence reads, using the method described in Skoglund et al. (30). |  |  |  |  |  |  |  |  |  |  |
| <sup>2</sup>Calibrated 95% range, SHcal13 (66, 67). |  |  |  |  |  |  |  |  |  |  |

To explore the population history of the Andean highlands, we first assess the genetic affinities of the prehistoric individuals and compare them to modern South Americans as well as other ancient Native Americans. Second, we construct a demographic model that estimates the timing of the lowland-highland population split as well as the population collapse following European contact, and finally, we explore evidence for genetic changes associated with selective pressures associated with the permanent occupation of the highlands, the intensification of tuber usage, and the impact of European-borne diseases.

## Results

### Samples and sequencing

We performed high-throughput shotgun sequencing of extracted DNA samples from seven individuals from three archaeological periods with proportions of endogenous DNA ranging from 2.60% to 44.8%. The samples span in age from 6,800 to 1,400 cal BP (Table S1). The samples from the Rio Uncallane site showed the highest endogenous content and five were chosen for frequency-based analyses and sequenced to an average read depth of 4.74x (Table 1). The oldest sample is from the SMP site, SMP5. The sample was directly dated to ~6,800 years BP and exhibited the highest endogenous DNA proportion among burials from the same site (3.15%) and was sequenced to an average read depth of 3.92x. The sample from the Kaillachuro site, at about 3,800 years BP (culturally dated), was sequenced to an average read depth of 1.44x. All samples exhibited low contamination rates (1-4%) and DNA damage patterns consistent with degraded DNA (Table S2, Fig. S2).

We also generated new data from two contemporary populations: 24 Aymara individuals from Ventilla, Bolivia, which is near Lake Titicaca (Fig. S1), were sequenced to an average read depth of 4.36x (Table S3), and an additional individual was sequenced to a read depth of 31.20x (*see SI Appendix*). The modern Huilliche-Pehuenche (n=39) from the south of Chile (Fig. S1) were genotyped on an Axiom LAT1 Array and imputed utilizing the 1,000 genome phase 3 (*17*) and the Aymara whole genome sequence data as a reference panel. Analysis using the program *ADMIXTURE* (*18*) revealed that all the modern samples had trivial amounts of non-indigenous ancestry (less than 5%).

### Genetic relationship between ancient and modern individuals

We employed outgroup *f*_3_ statistics to assess the shared genetic ancestry among the ancient individuals and 156 worldwide populations (*19*). C/T and G/A polymorphic sites were removed from the data set to guard against the most common forms of post-mortem DNA damage (*20*). Outgroup *f*_3_ statistics of a worldwide dataset demonstrate that all seven ancient individuals from all three time periods display greater affinity with Native American groups than with other worldwide populations (Fig. 1). Ranked outgroup *f*_3_ statistics suggest that the ancient individuals from all three time periods tend to share greatest affinity with Andean living groups (Fig. 1A).

**Figure 1.**
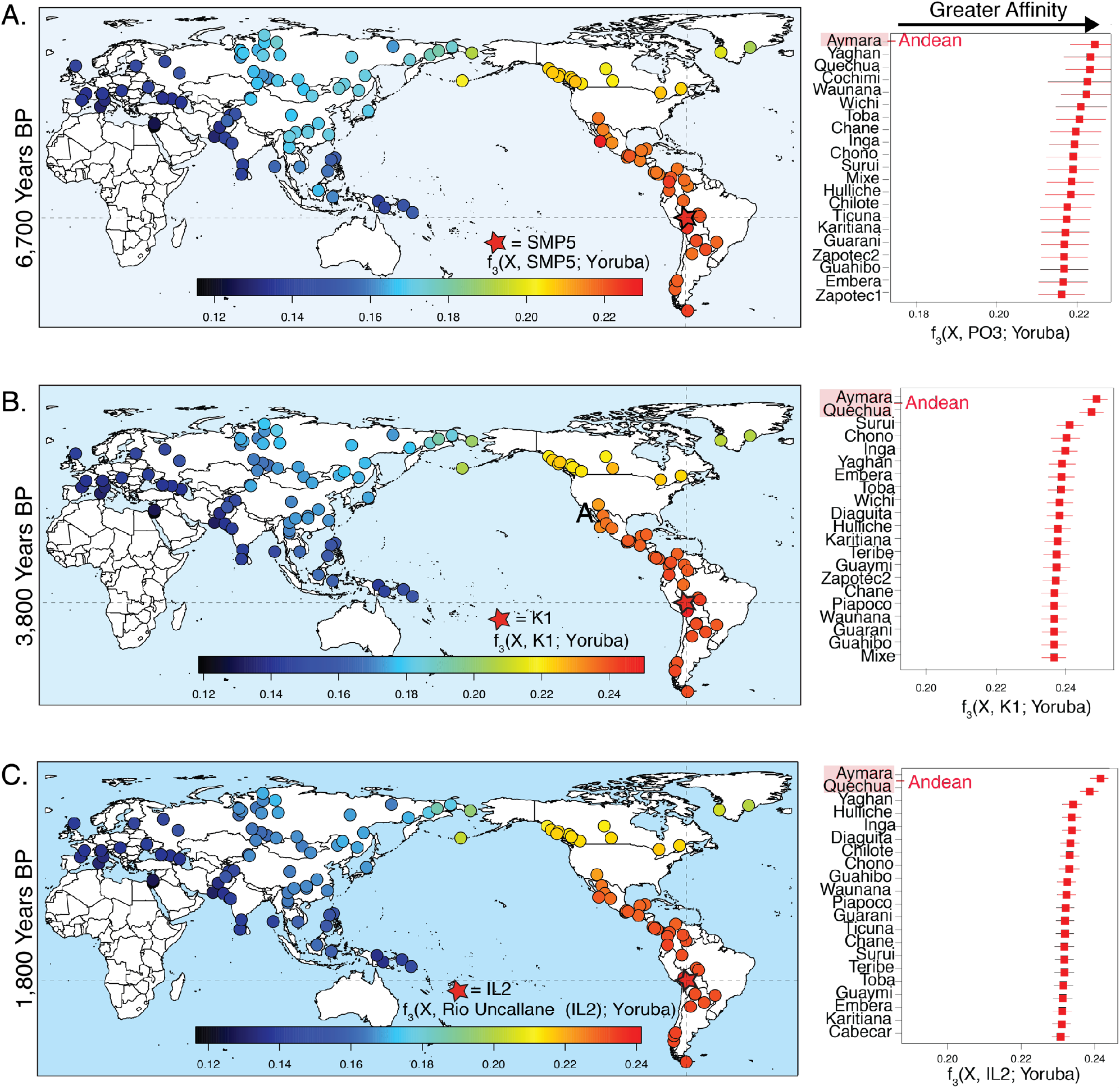
*f*_3_ statistics. (Left) Heat map represents the outgroup *f*_3_ statistics estimating the amount of shared genetic drift between the ancient Andean individuals and each of 156 contemporary populations since their divergence with the African Yoruban population. (Right) Ranked *f*_3_ statistics showing the greatest affinity of the ancient Andeans with respect to 45 indigenous populations of the Americas.

Principal components analysis reveals a tight clustering of the Rio Uncallane and Kaillachuro (K1) individuals, which overlays with modern Andean populations (Fig. 2). The oldest individual, SMP5, places closer to the intersect between the modern Andean groups of the Quechua and the Aymara. To further elucidate the relationship amongst the ancient Andean individuals and modern populations from South America, we examined maximum likelihood trees inferred with *TreeMix* (*21*). We observe that individuals from all three time periods form a sister group to the modern Andean high-altitude population, the Aymara (Bolivia) (Fig. 3A-C). The connection between the modern and ancient Andeans continues with the ADMIXTURE-based cluster analysis (Fig. 3D). Both the Rio Uncallane and K1 trace their genetic ancestry to a single component (shown in red), which is also shared by the modern Andean populations of the Quechua and the Aymara (Fig. 3D). SMP5 traces most of his ancestry to the same component, but also exhibits a component (shown in brown) found in Siberian populations, specifically the Yakut. USR1, a 12,000 year old individual from Alaska hypothesized to be part of the ancestral population related to all South American populations (*22*), also shares this component, as do previously published ancient individuals from North America (i.e., Anzick-1, Saqqaq and Kennewick).

**Figure 2.**
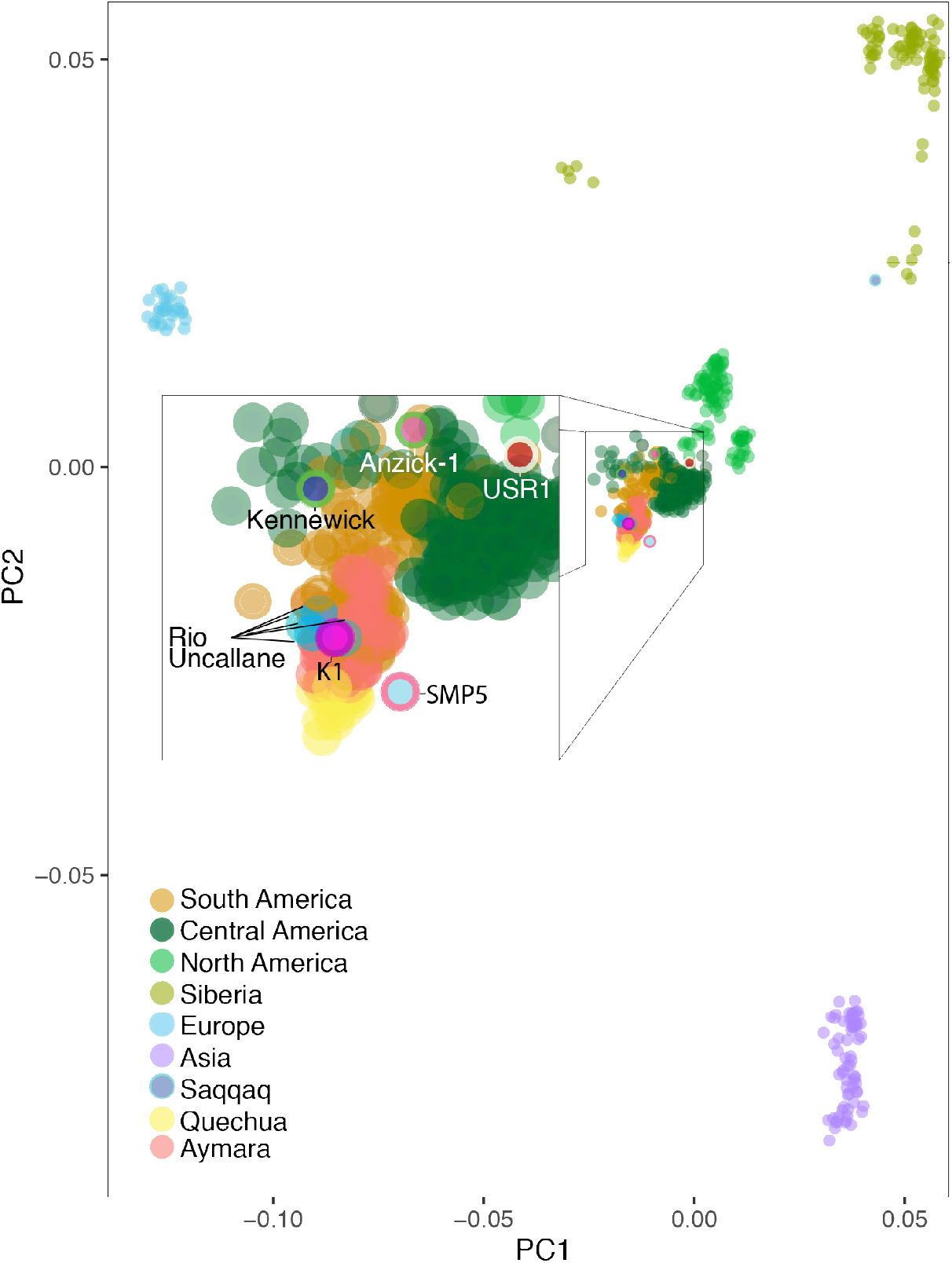
Principal components analysis. Principal components analysis projecting the ancient Andeans (IL2, IL3, IL4, IL5, IL7, K1, SMP5), USR1 (*22*) (Alaska), Anzick-1 (*62*) (Montana), Saqqaq (*63*) (Greenland), and Kennewick Man (*64*) (Washington) onto a set of non-African populations from Raghavan et al. (*19*), with Native American populations masked for nonnative ancestry.

**Figure 3.**
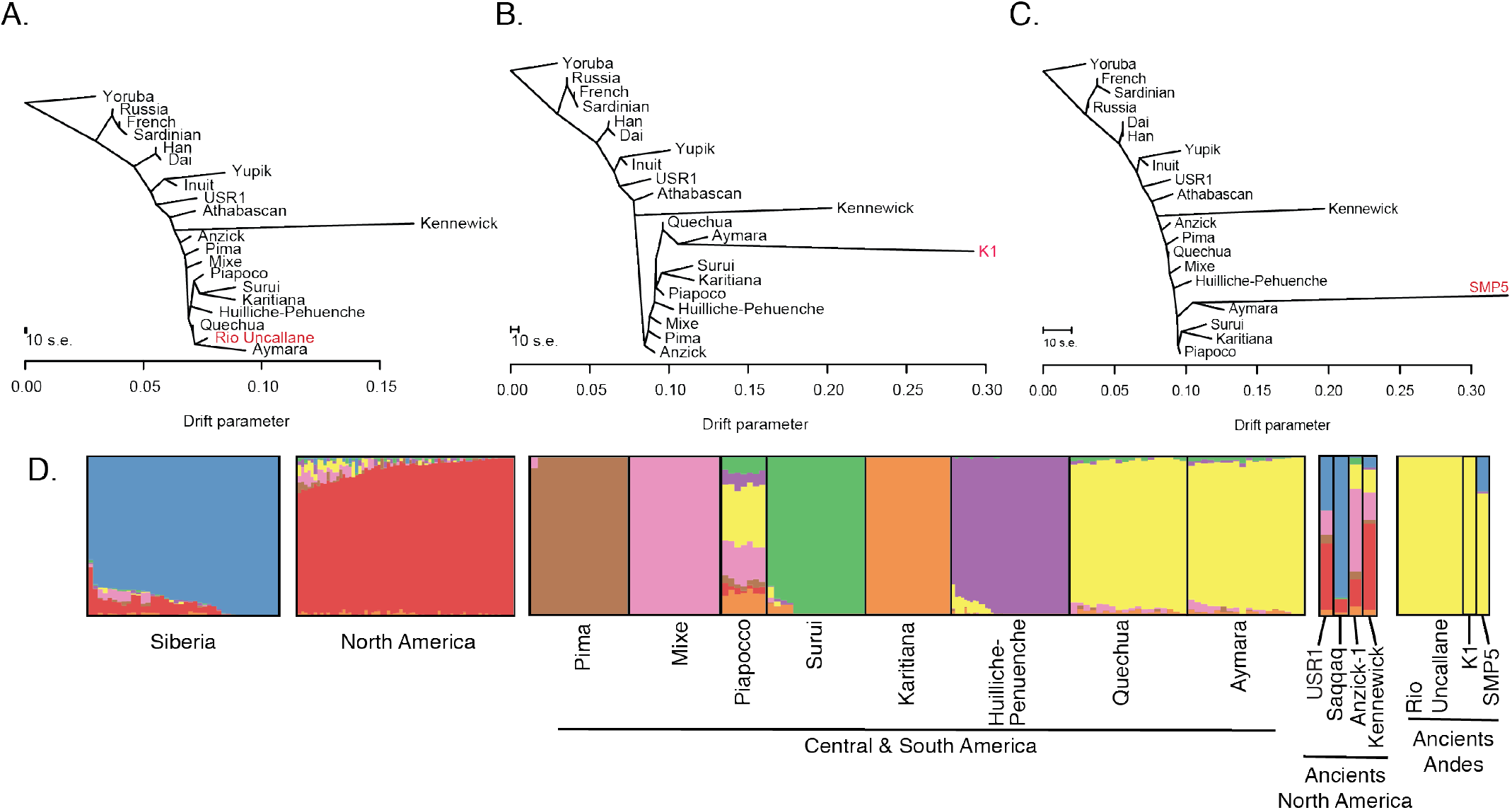
Genetic affinity of ancient Andeans to global and regional indigenous populations. (A-C) Maximum likelihood trees generated by *TreeMix* (*21*) using whole-genome sequencing data from the Simons Genome Project (*53*). (D) Cluster analysis generated by ADMIXTURE (*18*) for a set of indigenous populations from Siberia, the Americas, Anzick-1, Kennewick, Saqqaq, USR1, and the ancient Andeans. The number of displayed clusters is *K* = 8, which was found to have the best predictive accuracy given the lowest cross-validation index value. Native American populations masked for nonnative ancestry.

### Demographic model

Given that the above analyses provide an inference of genetic affinity in the Andes extending over roughly 4,000 years BP, and perhaps up to 7,000 years, we explored models relevant to the timing of the first permanent settlements in the Andes, as well as the extent of the population collapse occurring after European contact in the 1500s (Fig. 4). We utilized a composite likelihood method (Fastsimcoal2 (*23*)), which was informed by the site frequency spectra (SFS) of multiple populations, including the Rio Uncallane (n=5) (high altitude, pre-European contact), the Aymara (n=25) (high altitude, post-European contact), and the Huilliche-Pehuenche (n=39) (lowland population, post-European contact). We infer a split between low and high-altitude populations in the Andes occurring at 8,750 years (95% CI[8200, 9250]). We also infer the magnitude of the population collapse in the Andes after European contact to be a 27% reduction in effective population size and occurring 425 years ago (95% CI[400, 450]). The model also inferred the collapse experienced by the Mixe (included in the model for the split between Northern and Southern Hemisphere populations) and the Huilliche-Pehuenches after European contact, which were much more severe. We infer a 94% (95% CI[0.94-0.96]) collapse for the Mixe and a comparable 96% (95% CI[0.95-0.96]) collapse for the Huilliche-Pehuenche. The collapse difference between the high and low altitude populations from the Andes region is significant (Huilliche-Pehuenche vs Aymara: *P* < 0.0001 (95% CI[0.66-0.71]), chi-square test).

**Figure 4.**
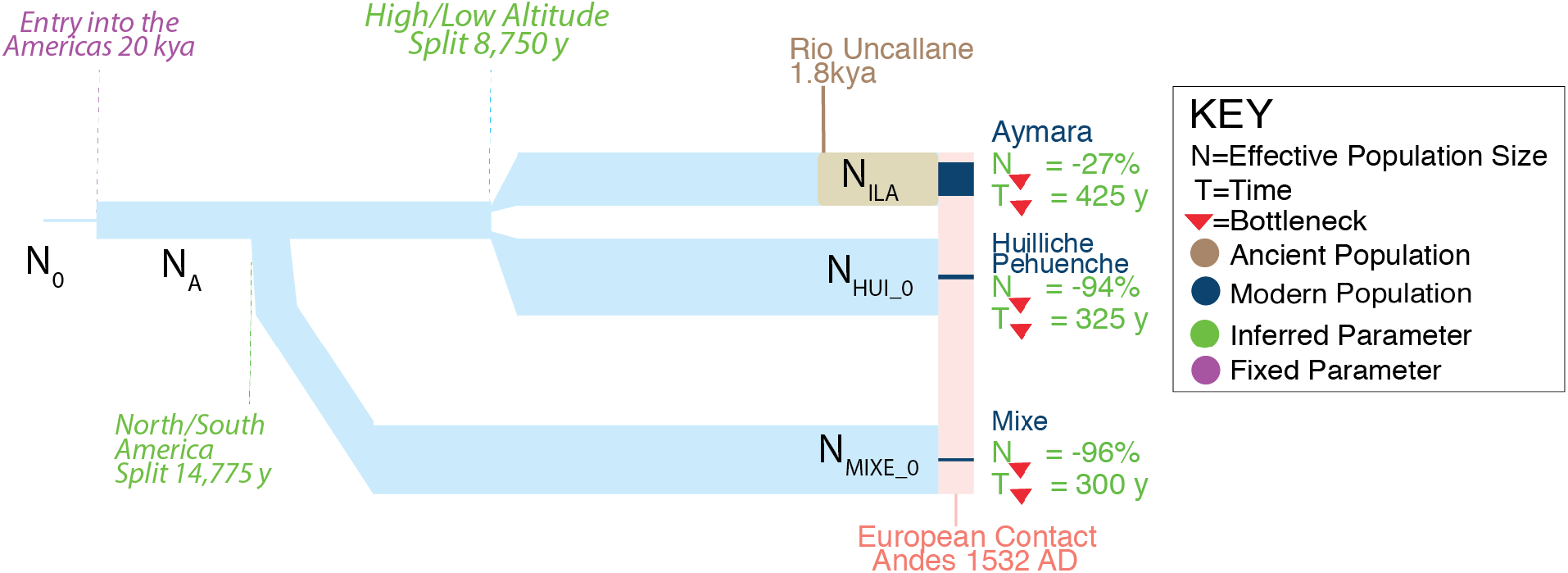
Demographic model of the Andes.

However, we were concerned with the heterogenous nature of the data used in the model, since the Huilliche-Pehuenche data was imputed from a SNP array. Therefore, we calculated heterozygosity on each population using an overlapping set of ascertained SNPs from the San. We found a similar pattern of effective population size changes, using heterozygosity as a proxy, as we did with the demographic model (Supplementary text, Fig. S3).

### Adaptive allele frequency divergence

Next, we turn to detecting the impact of natural selection. Permanent settlement of high altitude locales in the Andes likely required adaptations to the stress of hypobaric hypoxia, such as those detected in Tibetan, Nepali, and Ethiopian highland populations (*24–27*). Several genetic studies have also been performed on modern populations of the Andes (*25–31*). The availability of genetic variation data from an ancient Andean population, i.e., the Rio Uncallane samples, offers new opportunities for investigating adaptations in this region of the Americas. More specifically, the ancient data allows us to test for adaptations in a population that was not exposed to the confounding effects associated with the European contact, i.e., the demographic bottleneck and the exposure to additional strong selective pressures due to the introduction of new pathogens (*32, 33*). Therefore, we test for high allele frequency divergence resulting from adaptations to the Andean high altitude environment, utilizing a whole genome approach with the five Rio Uncallane individuals (~1800 BP), and we contrast them with a lowland population, the Huilliche-Pehuenche from Chile, who likely inhabited the region for several thousand years before the arrival of the Spanish (*34*). We utilized the Population Branch Statistic (PBS) (*27*) to scan for alleles that reached high frequency in the Rio Uncallane relative to the Huilliche-Pehuenche. The Han Chinese from the 1,000 Genomes Project (*17*) were utilized as the third comparative population. The single nucleotide polymorphisms (SNPs) showing the most extreme PBS score and, therefore, the highest differentiation in the ancient Andeans, corresponds to a region near *MGAM* (Fig. 5A, Table S4). *MGAM* is an intestinal enzyme associated with starch digestion (*35*) and could possibly be linked to the intensification of tuber use and ultimately the transition to agriculture (*8, 36*). The second strongest signal came from the gene *DST*, which encodes a cytoskeleton linker protein active in neural and muscle cells (*37*). *DST* has been associated with differential expression under hypoxic conditions (*38, 39*) and cardiovascular health (*40, 41*).

**Figure 5.**
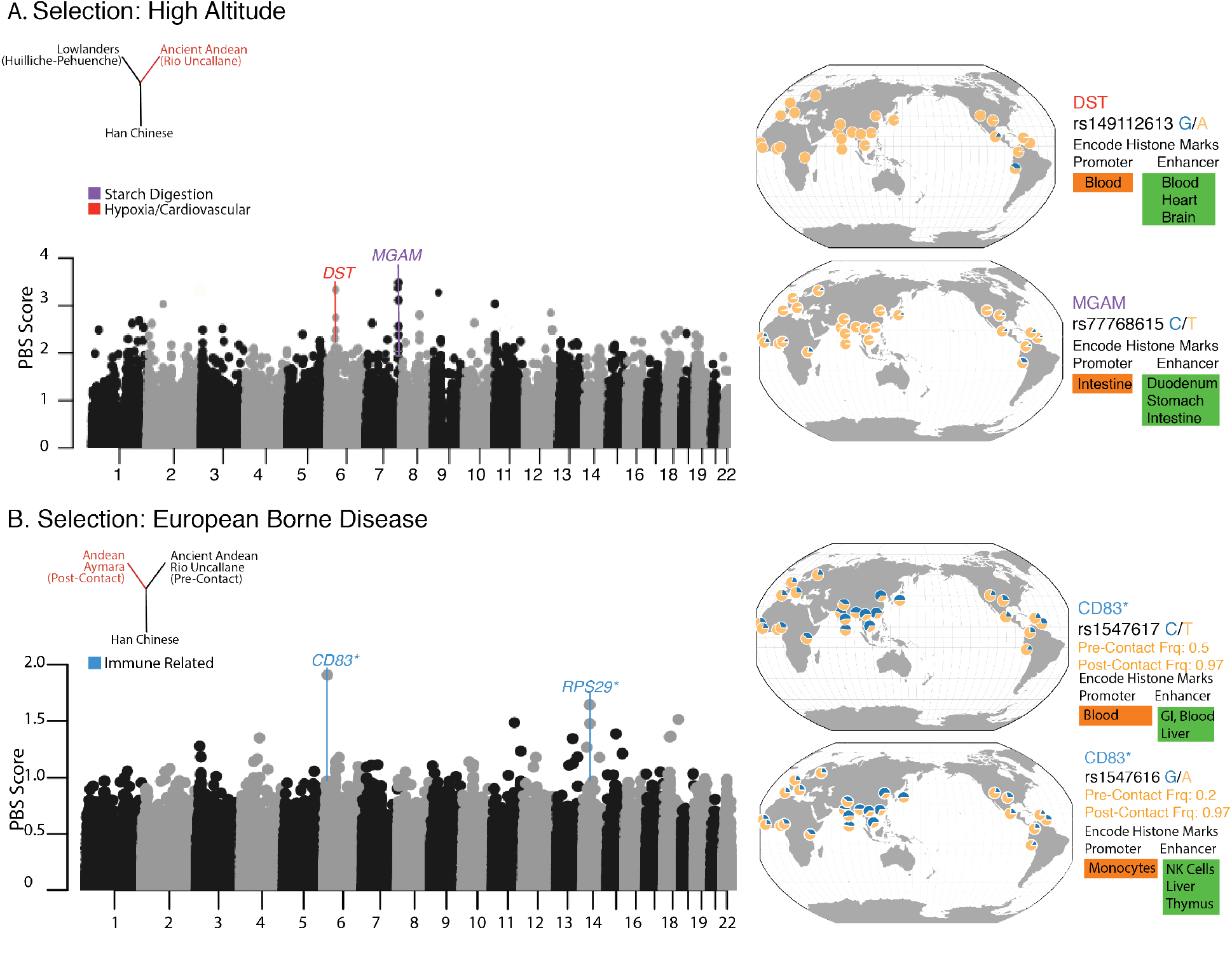
Selection scans. (A) PBS scans testing for differentiation along the Ancient Andean branch, with respect to the lowland population of the Huilliche-Pehuenche. (B) PBS scan testing for differentiation along the modern Aymara branch (post-European contact), with respect to the pre-contact ancient population of the Rio Uncallane. Selected global SNP frequencies (*65*) and histone modifications (*46*) are shown to the right, which are relevant to the strongest selection signals. *Indicates closest gene.

The second test for selection targeted hypotheses concerning the adaptation to European-introduced pathogens (Fig. 5B). The PBS was conducted with the Aymara (post-European contact), the Rio Uncallane (pre-European contact), and the Han Chinese (*17*). The strongest selection signal comes from SNPs in the vicinity of *CD83*, which encodes an immunoglobulin receptor and is involved in a variety of immune pathways, including T-cell receptor signaling (*42–44*). The gene has also been shown to be upregulated in response to the vaccinia virus infection (*45*), the virus used for smallpox vaccination. In addition, the differentiated SNPs associated with the *CD83* signal show a variety of chromatin alterations in immune cells, including monocytes, T-cells, and natural killer (NK) cells (*46*), suggesting that they may be functional. Lastly, a gene in the top 0.01% of the scan, *IL-36R*, codes for an interleukin receptor that has been associated with the skin inflammatory pathway and vaccinia infection (*47*).

## Discussion

South America is thought to have been populated relatively soon after the first human entry into the Americas, some 15,000 years ago (*3*). However, steep environmental gradients in western South America would have posed substantial challenges to population expansion. Among the harshest of these environments is the Andean highlands, which boasts frigid temperatures, low partial pressure of oxygen, and intense UV radiation. Despite this, however, humans eventually spread throughout the Andes and occupied them permanently. Archaeological evidence suggests that hunter-gathers entered the highlands as early as 12,000 years BP (*4*), with permanent occupation beginning around 9,000 years BP (*5–7*). The evidence presented here indicates genetic affinity between populations from different time periods in the high elevation Lake Titicaca region from at least 3,800 years BP, and possibly 7,000 years BP. This affinity extends to the present high-altitude Andean communities of the Aymara and Quechua.

Although our samples do not extend beyond 7,000 years BP, we were able to model the initial entry into the Andes after the split between North and South American groups. Our model, utilizing a 1.25 × 10^−8^ mutation rate, shows a correlation with archaeological evidence regarding the split between North and South groups occurring nearly 14,750 years ago (95% CI[14,225, 15,775]), which agrees with the oldest known site in South America of Monte Verde in southern Chile (~14,000 years BP) (*3*). The date for the split between low and high-altitude populations was inferred to 8,750 years (95% CI[8,200, 9,250]), which is younger than previously reported by a study utilizing modern genomes alone (*48*). This date provides a terminus ante quem time frame for the origins of adaptations known in modern highland populations.

We also present evidence for genes that may have been under selective pressure caused by environmental stressors in the Andes. Interestingly, none of our most extreme signals for positive selection were related to the hypoxia pathway. Instead, we find differentiated SNPs in the *DST* gene, which has been linked to the proper formation of cardiac muscle in mice (*40, 41*). Furthermore, the *DST* intronic SNP that was most differentiated (rs149112613) shows histone modifications associated with blood and the right ventricle of the heart (*46*). This correlates to Andean highlanders tending to have enlarged right ventricles associated with moderate pulmonary hypertension (*1*). This finding also parallels hypotheses proposed by Crawford et al. (*49*) that Andeans may have adapted to high altitude hypoxia via cardiovascular modifications.

The most extreme signal may represent adaptations to an agricultural subsistence and diet. The top ranked gene, *MGAM*, is associated with starch digestion (*50*). The associated high frequency SNPs in the ancient Andean population (Table S4), exhibit chromatin marks in cells from the gastrointestinal tract (Fig 5A). The variant may be highly differentiated between the ancient Andeans and the lowlanders (the Huilliche-Pehuenche) due to differences in subsistence strategies. The Huilliche-Pehuenche are traditionally hunter-gatherers, with archaeological evidence suggesting that their ancestors have been practicing this mode of subsistence for thousands of years in the region before European contact in the 1500s (*51*). In contrast, the Andes is one of the oldest New World centers for agriculture, which included starch-rich plants such as maize (~4000 years BP) (*52*) and the potato (~3400 Years BP) (*8*). Selection acting on the *MGAM* gene in the ancient Andeans may represent an adaptive response to greater reliance upon starchy domesticates. Recent archaeological findings based on dental wear patterns and microbotanical remains similarly suggest that intensive tuber processing and thus selective pressures for enhanced starch digestion began at least 7000 years ago (*8, 36*). Furthermore, we see a similar signal (top 0.01%) when we contrast the hunter-gatherers from Brazil (Karitiana/Surui, sequence data (*53*)) with the ancient Andeans, as well as with the Aymara vs. the Huilliche-Pehuenche and the Karitiana/Surui. One further note, we did not detect *amylase* high copy number in the ancient Andes population before European contact, suggesting a different evolutionary path for starch digestion in the Andes when contrasted with Europeans (*54*).

Selection with respect to the environment in the Andes is not limited to the ancient past. In 1532, the environment radically changed with the arrival of the Spanish (*55*). Not only were long-standing states and social organizations disrupted, but the environment itself was altered with the arrival of European-introduced pathogens, which may have preceded the arrival of the Spanish via trade routes (*55*). Some of the most devastating epidemics were related to smallpox, occurring in the 1500s and 1600s (*56*). These combined factors are thought to have decimated the local populations (*56*). We infer the population decline in the Andes, utilizing the ancient and modern Andeans, and found the decline in effective population size to be 27% (95% CI[0.23, 0.34]). This is a modest decline compared to archaeological and historical estimates, which reached upwards of 90% of the total population (*13*). In contrast, the model infers a much more severe collapse for lowland Andean populations of the Huilliche-Pehuenche, exceeding 90%. We also explored alternative scenarios to make sure the model was not biasing the inferred collapse and found that the estimated population size reduction remained significantly less severe in the highland compared to the lowland populations (see Supplement). Although we did not have pre-Contact ancient samples for these populations in Chile to inform the model, the large difference suggests that high-altitude populations may have suffered a less intense decline compared to the more easily accessible populations near sea level in the Andes region. This is also supported by long-lasting warfare in the Chilean lowland region with the Spanish that lasted well into the 19^th^ Century (*55*).

Our data also show that the populations in the modern Andes have high genetic affinity with the ancient populations preceding European contact. Although a strict continuity test (*57*) that does not allow for recent gene flow was not significant (see Supplement), modern Andeans are likely the descendants of the people that suffered the epidemics described in historical texts. In the Andes, missionary reports suggest that disease may have arrived before formal Spanish contact in 1532 (*58*) and that the first epidemics were likely caused by smallpox (*55*). We infer that selection acted within the past 500 years on the immune response making it likely that modern Andeans descend from the survivors of these epidemics. The selection scan along the branch of the modern Andeans, contrasted with the ancient group, revealed the strongest signal to be associated with an immune gene connected to smallpox, *CD83* (*45*). The second most highly differentiated SNPs were in the vicinity of *RPS29*, which codes for a ribosomal protein, and is involved in viral mRNA translation and metabolism, including that of influenza (*59*). Another top gene, *IL-36R* (rank #36), is thought to have evolved alternative cytokine signaling to compensate for viruses, such as pox viruses, that can evade the immune system (*60*). Furthermore, the top SNP associated with *IL-36R* (rs1117797) exhibits a QTL signal associated with *IL18RAP*, a gene involved in mediating the immune response to the vaccinia virus (*61*). The relative strength of these signals and the role played by the associated genes, may indicate that selection favored alleles that directly affected the pathogenicity of the diseases encountered by the ancestors of the epidemic survivors.

In conclusion, human adaptation to the Andean highlands involved a variety of factors and was complicated by the arrival of Europeans and the drastic changes that followed. Despite harsh environmental factors, the Andes were populated relatively early after entry into the continent. The adaptive traits necessary for permanent occupation may have been selected for in a relatively short amount of time, on the order of a few thousand years. Given the multifaceted nature of the adaptation, we are not surprised to find genetic affinity in the populations of the Andes dating to at least 4,000 years BP, and possibly extending to 7,000 years BP.

## Acknowledgments

We like to thank John and Sarah Blangero for their contribution with the Aymara samples. We would like to thank Choongwon Jeong and Shigeki Nakagome for helpful discussions throughout the project. We would like to thank Mike DeGiorgio for his help with the F3 statistic analysis. Field support was provided by Collasuyo Archaeological Research Institute, Cecilia Justo Chavez, Virginia Incacoña Huaraya, Mateo Incacoña Huaraya, Nestor Condori Flores, Albino Pilco Quispe, Daniel Pilco Incacoña, Dario Pilco Incacoña, Karen Pilco Incacoña, Luz Mery Pilco Incacoña, Lauren Hayes and the community of Mulla Fasiri, Peru.

## Data and materials availability

Ancient and modern DNA sequences are available from NCBI Sequence Read Archive, accession no. PRJNA470966. The Huilliche-Pehuenche SNP data will be available via a data access agreement with Ricardo Verdugo at the Universidad de Chile.

## Permits

Archaeological data recovery at SMP and international export of artifacts were carried out under Peruvian Ministry of Culture permit no. 064-2013-DGPA-VMPCIC/MC and 138-2015-VMPCIC/MC. Multiple permits were issued by the Peruvian Instituto Nacional de Cultura for research at Kaillachuro and the Rio Uncallane sites from 1994-2002.

## Supplementary Materials

Materials and Methods

Figures S1-S5

Tables S1-S6

References (*59–93*)

